# A Novel Slow-Progressive Knee Osteoarthritis Murine Model Induced by Non-Invasive Posterior Cruciate Ligament Rupture in Mice

**DOI:** 10.64898/2026.05.11.724206

**Authors:** Saaya Enomoto, Kohei Arakawa, Kei Takahata, Michiaki Sato, Himari Miyamoto, Riku Saito, Yuna Usami, Koyo Nogi, Takanori Kokubun

**Author notes:** **Full contact details of the co-corresponding author:** Takanori Kokubun, Ph.D., Graduate School of Health, Medicine, and Welfare, Saitama Prefectural University, 820 Sannomiya, Koshigaya, Saitama, 343-8540, Japan. **Author email addresses:** S. Enomoto;, K. Arakawa;, K.Takahata;, M. Sato;, H. Miyamoto;, R. Saito;, Y. Usami;, K. Nogi. **Author information:** Saaya Enomoto and Kohei Arakawa contributed equally to this work.

## Abstract

**Objective:** Recently, alternatives to animal testing, such as new approach methodologies, are being developed in the orthopedic research field; animal models still provide valuable insights into the pathogenesis of knee osteoarthritis (OA). However, commonly used models develop OA much more rapidly and severely than those observed in human patients. We aimed to develop a novel murine model that closely mimics the slow progression of human OA with posterior Cruciate ligament (PCL) rupture.

**Design:** 12-week-old C57BL/6 mice were induced to PCL-rupture (PCL-R) by manually applying an external tibial posterior translation force. We analyzed joint kinematics, histological observations, and bone structure to confirm the absence of concurrent injury on day 0. Then, joint stability and the pathophysiological progression of knee OA were analyzed at 8, 16, and 34 weeks post-PCL-R. The destabilized medial meniscus (DMM) model was also analyzed to compare the OA progression.

**Results:** Non-invasive PCL-R intervention induced the complete rupture in the central region of PCL without concurrent injury. The PCL-R group showed larger posterior tibial deviation than the INTACT (*P*=0.008). Regarding the range of motion in the PCL-R group, there was no limitation in range of motion on day 0, but extension limitations occurred at weeks 16 and 34 weeks. Histologically, articular cartilage degeneration in PCL-R was milder than DMM. In the subchondral bone, micro-CT reconstruction images indicated that, compared with the INTACT group, the DMM group observed progressive subchondral bone formation from 16 weeks post-surgery. In contrast, the PCLR group maintained the subchondral bone structure even at 34 weeks.

**Conclusions:** PCL-R model induced mild abnormal mechanical stress depending on posterior instability, and cartilage degeneration occurred more slowly in this model than in DMM models.

## INTRODUCTION

Knee osteoarthritis (OA) is a slowly progressive musculoskeletal disease characterized by fibrillation and degeneration of articular cartilage. Knee OA can be divided into two subtypes based on onset mechanism: primary and secondary^1,2^. Primary knee OA gradually degenerates the cartilage and joint tissues without any apparent underlying causes, such as a fracture or surrounding tissue injury. Secondary knee OA is the consequence of either an abnormal mechanical force, such as compression or shear force on the cartilage underlying the post-traumatic (PT) injury due to a ligament or meniscus injury, or abnormal articular homeostasis, such as rheumatoid arthritis^3^. The percentage of secondary knee OA to total knee OA is estimated to be 12-42%^4^. Knee OA is the leading cause of joint disability worldwide, resulting in a very high medical and socioeconomic burden^5^. In addition, OA patients have been reported to have a four times increase in healthcare costs compared to those without OA, and the burden of OA is continuously increasing in many countries^6,7^. In response to this issue, treatment for knee OA has focused on improving various symptoms, especially joint pain. However, the prevalence of OA is increasing rapidly^8,9^, and the strategy for OA needs to shift from disease treatment to prevention.

To treat and prevent knee osteoarthritis, some animal models were used to analyze the biological mechanisms underlying cartilage degeneration. The advantage of the surgical model over the spontaneous model is that it has less variability and can be analyzed quickly with fewer mice^10^. A representative animal model of OA is the Anterior Cruciate Ligament Transection (ACL-T) model^11,12^, which has been used primarily in rodents and across various animal species. The ACL-T model induces knee joint instability, and cartilage degeneration progresses quickly^13^. Then, the destabilization of the medial meniscus (DMM) model was reported as a relatively reproducible and slowly progressive model ^14^. However, the commonly used ACL-T and DMM models showed a comparatively severe pathological state within a shorter period than those for human OA patients. These models could be applicable to analyze the degradation process of articular cartilage from mild to severe, and they are a valuable preclinical model for investigating the effects of treatments and regenerative medicine in degenerated cartilage. Simultaneously, these rapid models are not suitable for establishing a prevention strategy for knee OA, as they do not capture the intra-articular pre-OA condition or the mechanism of the initial joint inflammation.

Recently, new approach methodologies (NAMs) that alternate to animal testing, such as organ-on-a-chip devices and micro-physiological systems, are being developed in the orthopedic research field^15,16^. Nevertheless, animal models still provide valuable insights into understanding the pathogenesis of knee OA. However, a novel knee OA model is needed to explore the pre-intra-articular condition and onset mechanism of primary knee OA. A novel model should have several key features: gradual cartilage degeneration over a longer time course, continuous joint instability, and no direct effect on the intervention itself. Here, we developed a novel long-term progressive knee OA rodent model with non-invasive rupture of the Posterior Cruciate ligament (PCL). PCL is one of the supporting tissues of the knee joint stability. The PCL is known to be a thicker and stronger ligament than the ACL^17^, which controls posterior tibial translation. In contrast to ACL injuries, OA changes after PCL injury are less severe in human patients.^8^ This characteristic difference is thought to be due to the less instability that occurs in PCL damage than in ACL damage ^17^. Clinically, PCL dysfunction induces malalignment between the tibia and femur, but to a lesser extent than that of other knee ligaments in patients ^18^. We hypothesized that the PCL-R model would show slower OA progression than current joint instability-induced knee OA models. Additionally, surgical intervention also affected the biological response in the knee intra-articular environment. A non-invasive ACL-R model has been reported by Christiansen et al. ^19^, and we have further established a novel non-invasive ACL-R mouse model without intra-articular and subchondral tissue damage ^20^. In this study, we applied these non-invasive methods to create a PCL-R model. The purpose of this study was to clarify the characteristics and phenotypes of the novel non-invasive PCL-R model.

## MATERIALS AND METHODS

### Animals and experimental design

This study was approved by the Animal Research Committee of Saitama Prefectural University (approval number: 2021-11). The animals were handled in accordance with relevant legislation and institutional guidelines for humane animal care. Here, 76 C57BL/6 male mice were used. 30 mice were intervened with PCL-R on the left hindlimb, and the contralateral limb was used as INTACT. All mice were housed in plastic cages under a 12-hour light/dark cycle. The mice were allowed unrestricted movement within the cage and had free access to food and water. The mice were sacrificed at 0 day, 8, 16, and 34 weeks (n=10 each), and knee joints were harvested for each analysis. At each time point, 5 mice were used for radiographic and histological analysis, and the remaining 5 for microCT analysis. To compare the PCL-R model with existing current knee OA models, we created the DMM model. We divided 30 male mice into the DMM group. The DMM group was sacrificed at 8, 16, and 34 weeks (n=10 each) for histological and micro CT analysis.

### PCL-R intervention procedures

All procedures were performed on the left knee joint of each mouse under a combination of anesthetic drugs (medetomidine, 0.375 mg/kg; midazolam, 2.0 mg/kg; and butorphanol, 2.5 mg/kg). The hip and knee joints were fixed at 90° on a stand. The tibial plateau was manually pushed vertically, causing PCL rupture due to the relative posterior dislocation of the tibia (Fig.2A).

**Figure 1.**
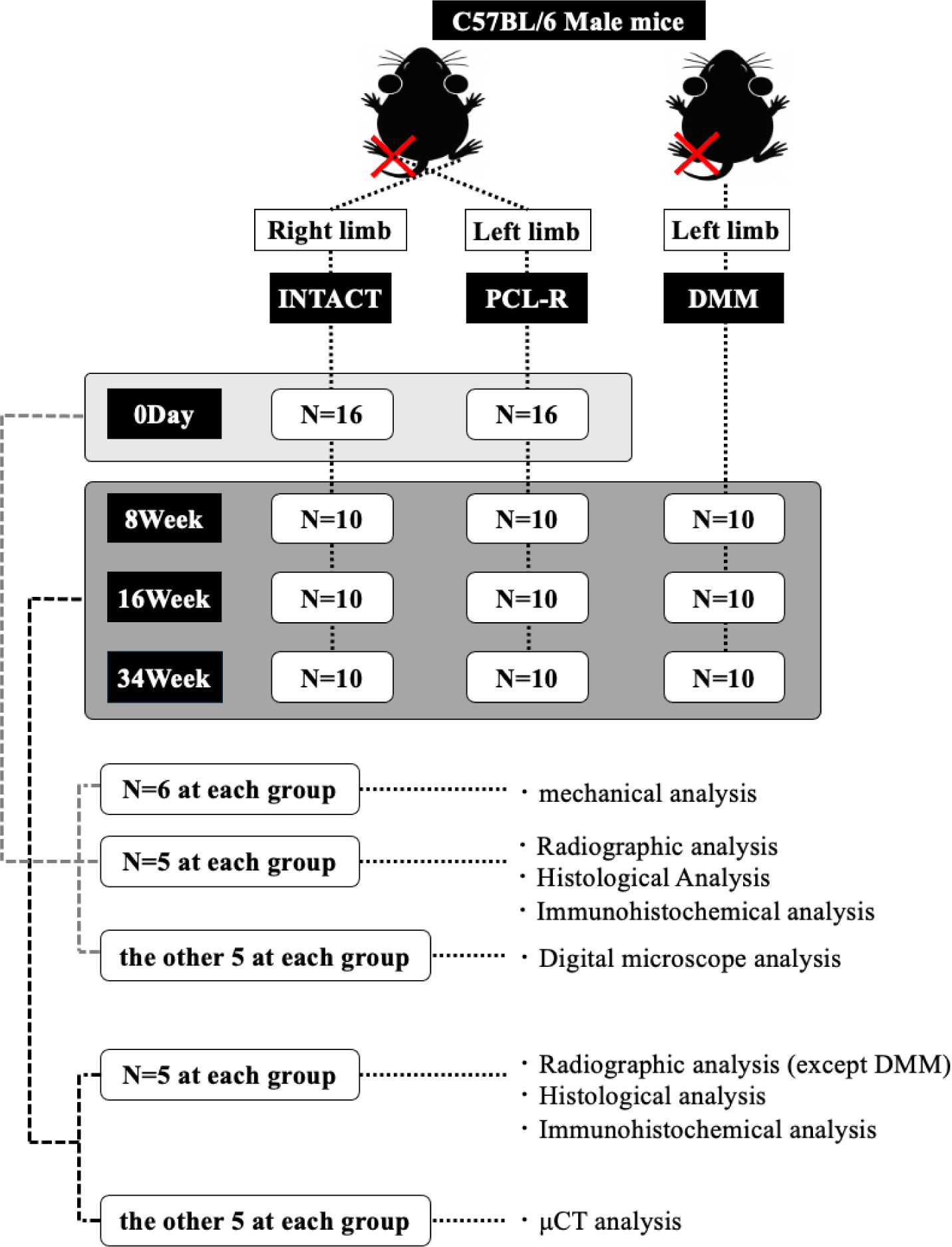
Experimental design. We created the non-Invasive PCL-R model and performed a mechanical, radiographic, histological, and macroscopical analyses on the same day to confirmed that the PCL was ruptured and that there was no intra-articular injury after intervention. In addition, we performed radiographic, histological and μCT analyses at 8, 16, and 34 weeks after the PCL-R and DMM intervention to compare the cartilage degeneration and subchondral bone change among INTACT, PCL-R, and DMM.

**Figure 2.**
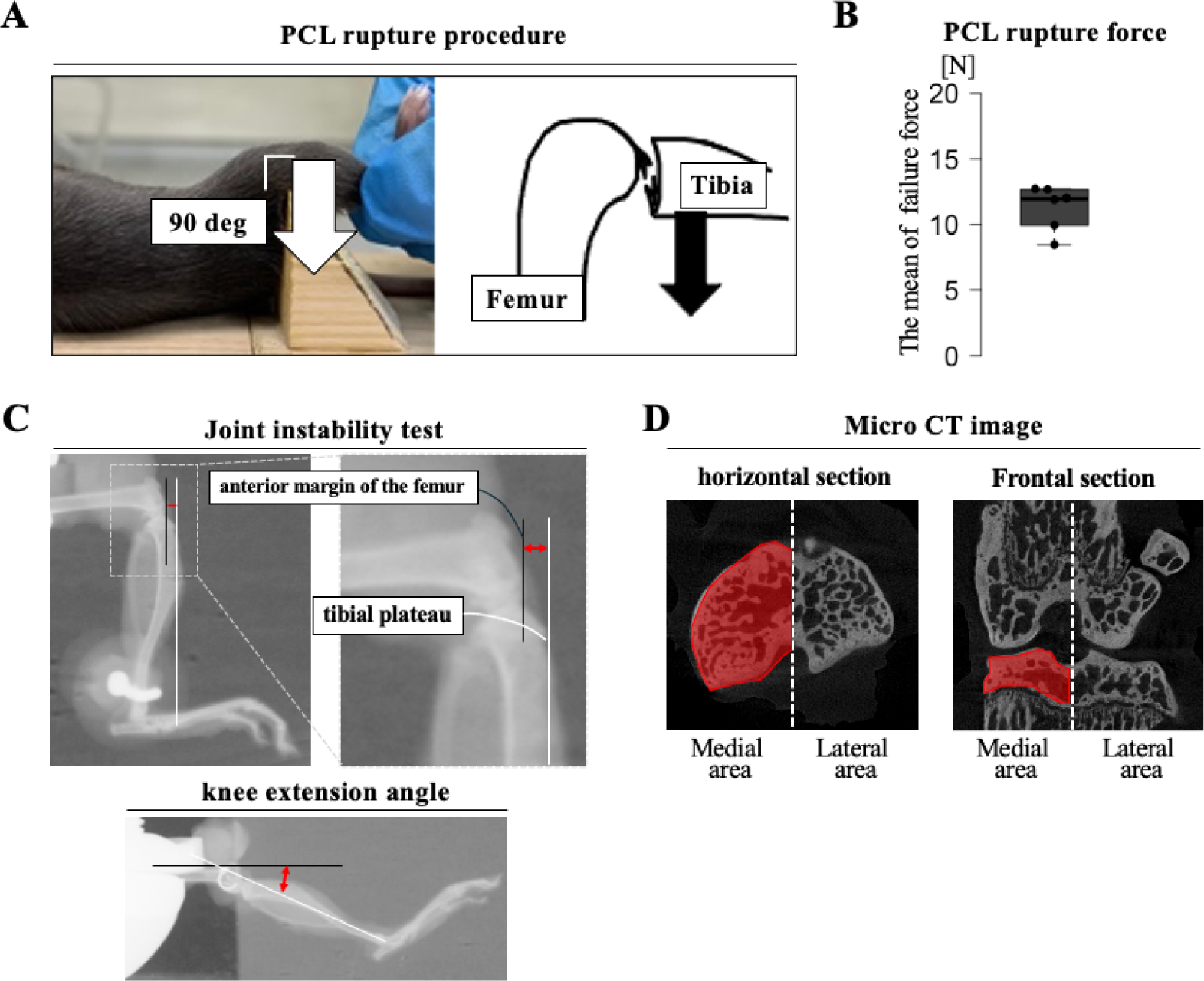
The intervention procedures of the non-invasive PCL-R model. (A) The mice was laid on its back with their hip and knee joints fixed at 90 degrees. After that, the tibia plateau were pushed manually. (B) The result of the PCL rupture force. The mean failure force causing PCL rupture ranged between 8.56 and 12.7 N. (C) The methods of joint instability test and knee extension angle evaluation. Joint instability was measured as the distance between the anterior margin of the femur and the tibial tuberosity. The knee extension angle was measured as the angle formed between the femoral longitudinal axis and the line connecting the tibial tuberosity and the anterior margin of the ankle joint. (D) Region of interest (ROI) in micro-CT analysis. ROI was defined as the entire medial tibial subchondral bone.

### Measurement of the PCL rupture force

Before the experiment, 6 mice were used for the assessment of PCL rupture force following the method established by Takahata et al.^20^ (Fig.2B). The MDB-25 load cell (Transducer Techniques, CA, USA), along with a load cell amplifier (HX711, SparkFun, CO, USA), was employed to determine the PCL rupture force. Calibration of the load cell was first performed using a 30.7271 g weight for offset adjustment, from which a correction function was obtained, and voltage values were calibrated accordingly. After calibration, the load cell amplifier was connected to a PC, and data acquisition was performed using an Arduino and Jupyter Lab. The mouse femoral condyle was then loaded along its longitudinal axis using a bolt attached to the load cell. The mean failure force and the loading rate were calculated from the mechanical data recorded by the load cell.

### Radiographic analysis

#### □. Joint movement test

At each time point, mice were sacrificed, and the knee joints were harvested. Prior to histological analysis, the knee joints were imaged by X-ray. After that, we performed anterior and posterior drawer tests to evaluate anteroposterior joint deviation using the method described in Arakawa et al. Briefly, the knee joints were fixed at 90° of flexion in the device, and the proximal tibia was pulled forward or backward with a constant-force spring (0.05 kgf; Sanko Spring Co., Ltd., Fukuoka, Japan). X-ray images were then taken, and the anterior or posterior deviation was measured as the linear distance between a perpendicular line drawn from the anterior end of the femoral condyle and the tibial tuberosity (ImageJ; National Institutes of Health, Bethesda, MD, USA). The assessment was conducted by two independent observers (SR and MH) who were blinded to all other sample information. For the analysis, we measured the distance between the anterior margin of the femur and the tibial tuberosity. The mean of the two observers’ scores was used as the representative value. (Fig.2C)

#### □. Evaluation of knee extension angle

The distal end of the tibia was pulled in the direction of extension by pulling obliquely upward with a constant force spring (0.05 kgf; Sanko Spring Co., Fukuoka, Japan). It was manually confirmed that the extension could go no further. X-ray images were then taken, and the angle formed by the long axis of the tibia and femur was calculated. The assessment was conducted by two independent observers (SR and MH) who were blinded to all other sample information. For the analysis, we measured the angle formed between the femoral longitudinal axis and the line connecting the tibial tuberosity to the anterior margin of the ankle joint. The mean of the two observers’ scores was used as the representative value. (Fig.2C)

### Histological analysis

The harvested knee joints were fixed with 4% paraformaldehyde for 2 days, decalcified in 10% ethylenediaminetetraacetic acid for 2 weeks, dehydrated, and embedded in paraffin. The samples were cut in the sagittal plane (6 μm thickness) using a microtome. To confirm whether the PCL was completely ruptured, we performed microscopic observations using HE staining. In addition, to evaluate articular cartilage degeneration, we performed Safranin-O/fast green staining. At day 0, we confirm that the PCL-R intervention doesn’t cause medial and lateral intra-articular injury. The Osteoarthritis Research Society International (OARSI) histopathological grading system was used to assess cartilage degeneration^21^ by two independent observers (KA and KT) blinded to all other sample information. We used the center of the medial tibial plateau for analysis. The mean of the two observers’ scores was used as the representative value.

### Immunohistochemical analysis

To assess the expression of type X collagen (Col X), immunohistochemical staining was performed using the avidin-biotinylated enzyme complex method and a Vectastain Elite ABC Rabbit IgG Kit (Vector Laboratories, Burlingame, CA, USA). The tissue sections were deparaffinized with xylene and ethanol, and antigen activation was performed using proteinase K (Worthington Biochemical Co., Lakewood, NJ, USA) for 15 min. Then, endogenous peroxidase was inactivated with 0.3% hydrogen peroxide/methanol for 30 minutes. Nonspecific binding of the primary antibody was blocked using normal goat serum for 30 min. The sections were then incubated overnight at 4℃ with anti-Col X antibody (1:100, NBP3-03757, Novus Biologicals). Afterward, the sections were incubated with a biotinylated secondary anti-rabbit IgG antibody and stained with the Dako Liquid DAB Substrate Chromogen System (Dako, Glostrup, Denmark). The cell nuclei were stained with 25% hematoxylin. A region of interest (ROI) of 40,000 µm² (200 µm × 200 µm) was defined in the contact area of the knee joint fixed at 90° flexion. The ratio of Col X–positive cells to chondrocytes was then calculated.

### Subchondral bone structural analysis using micro-computed tomography (micro-CT)

The harvested knee joints were fixed with 4% paraformaldehyde for 2 days. The knee joints were scanned using a micro-CT system (Skyscan 1272, BRUKER, MA, USA) with the following parameters: pixel size, 6 μm; voltage, 60 kV; current, 165 μA. The reconstructed image was acquired using the NRecon software (BRUKER, MA, USA). We set the entire medial tibial subchondral bone as ROI (Fig.2D). We then calculated the bone volume/tissue volume fraction (BV/TV, %), trabecular thickness (Tb.Th, mm), trabecular number (Tb.N, 1/mm), and trabecular separation (Tb.Sp, mm) using CTAn software (BRUKER, MA, USA).

### Roughness evaluation of cartilage surface layer (digital microscope)

The samples slaughtered immediately after the PCL-R intervention and used for CT analysis were evaluated for cartilage surface roughness after imaging. The knee joint was dissected to expose the tibial articular surface by removing the femur, followed by removal of the meniscus, muscles, and ligaments, and imaging with a digital microscope (VR-6000, Keyence, Osaka, Japan). After setting the photographed articular surface as the reference plane, cartilage surface roughness analysis was performed for each of the medial and lateral sides of the tibia. The ROI was defined as an ellipse including the area not covered by the meniscus. The proportion of the ROI was then calculated by dividing its area by the total area of the medial and lateral articular surfaces (minimum 23%, maximum 36%).

### Statistical analysis

Statistical analysis of the measured data was performed using R software version 3.6.1. First, the normality of all data was verified using the Shapiro–Wilk test. The Wilcoxon rank-sum test was performed for the nonparametric comparison of two groups in the radiographic analysis and cartilage roughness evaluation. Histological, immunohistochemical, and subchondral bone analyses were compared among three groups using the nonparametric Kruskal–Wallis test. Post-hoc comparisons were subsequently performed using the Steel–Dwass test. Data are presented as medians with interquartile ranges, and statistical significance was set at P < 0.05.

## RESULTS

### Measurement of the PCL rupture force

We used a load cell to measure the rupture force during PCL-R modeling. As a result, the rupture strength ranged between 8.56 to 12.7 N (Fig.2B).

### No soft tissue injuries, except for PCL rupture, were detected immediately after intervention

X-ray images and graphs showing the amounts of anterior and posterior deviation, and the extension angle were shown in Fig.3 (A). The posterior deviation in the PCL-R group was significantly higher than that in the INTACT group (*p*=0.008). There was no significant difference in the anterior deviation between the PCL-R group and the INTACT group (*p*=0.69). There was no significant difference in the knee extension angle between the PCL-R group and the INTACT group (*p*=0.42). The ruptured PCL image with the H&E staining and the medial and lateral knee joints with the Safranin O Fastgreen stain are shown in Fig.3 (B,C). We confirmed complete PCL rupture in all PCL-R mice (Fig.3B). However, we didn’t find cartilage and meniscus damage in both medial and lateral knee joints in all PCL-R mice (Fig.3C). The results of the cartilage surface roughness analysis are shown in Fig.3 (D). There was no significant difference in the cartilage roughness between the PCL-R group and the INTACT group (medial area; *p*=0.49, lateral area; *p*=0.34).

**Figure 3.**
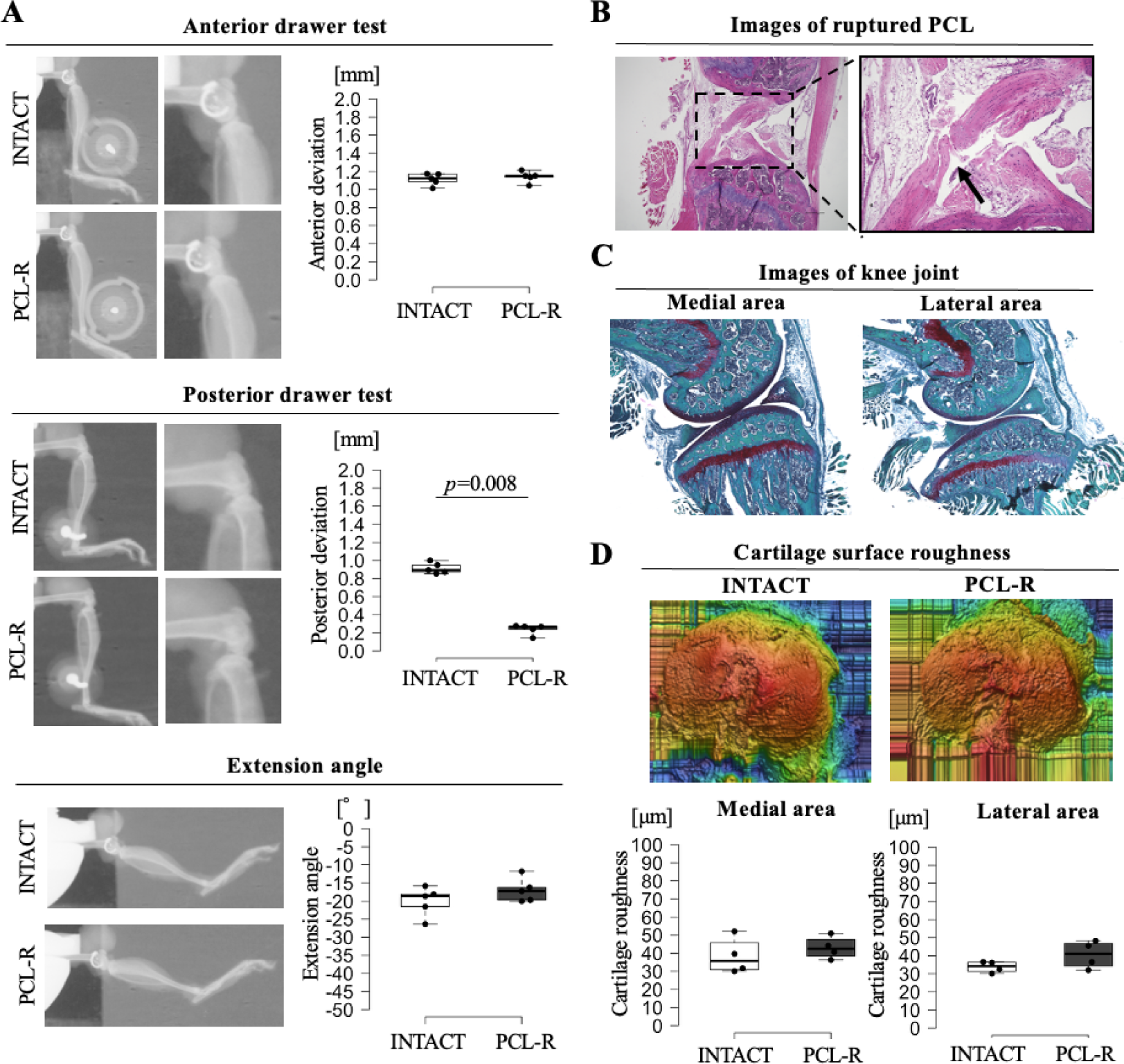
Evaluation of PCL rupture and cartilage surface roughness. (A) Evaluation of PCL rupture using a soft X-ray device. In the anterior tibial deviation, there were no significant difference between PCL-R and INTACT. The posterior tibial deviation was significantly increased in the non-invasive PCL-R group compared to that of the INTACT group (*p*=0.008). The data are presented as the median with interquartile range. Results of the extension angle test. There were no significant difference between PCL-R and INTACT. The data are presented as the median with interquartile range. (B-C) Histological analysis of the PCL and intra-articular tissues using H&E and safranin-O/fast green staining. In the PCL-R group, completely ruptured PCL was confirmed. however, no articular cartilage or meniscus injuries were observed. All scale bar: 300 mm. (D) The roughness of the cartilage surface. There are no difference between two groups. The data are presented as the median with interquartile range.

### Assessment of knee joint kinematic characteristics in the long term

To evaluate the validity of each model, we performed an anterior and posterior drawer test (Fig.4A). At all timepoints, the posterior deviation in the PCL-R group was significantly higher than that in the INTACT group (*p*=0.008 at all timepoints). On the other hand, there was no significant difference in the anterior deviation between the PCL-R group and the INTACT group (8 weeks; *p*=0.58, 16 weeks; *p*=0.69, 34 weeks; *p*=0.42). The knee extension angle was measured to assess the effect of changing the contact area (Fig.4B). At 8 weeks, there was no significant difference in the knee extension angle between the PCL-R group and the INTACT group (*p*=1.00). At 16 and 34 weeks, the knee extension angle in the PCL-R group was significantly lower than that in the INTACT group. (at 16 weeks; *p*=0.008, at 34 weeks; *p*=0.016).

**Figure 4.**
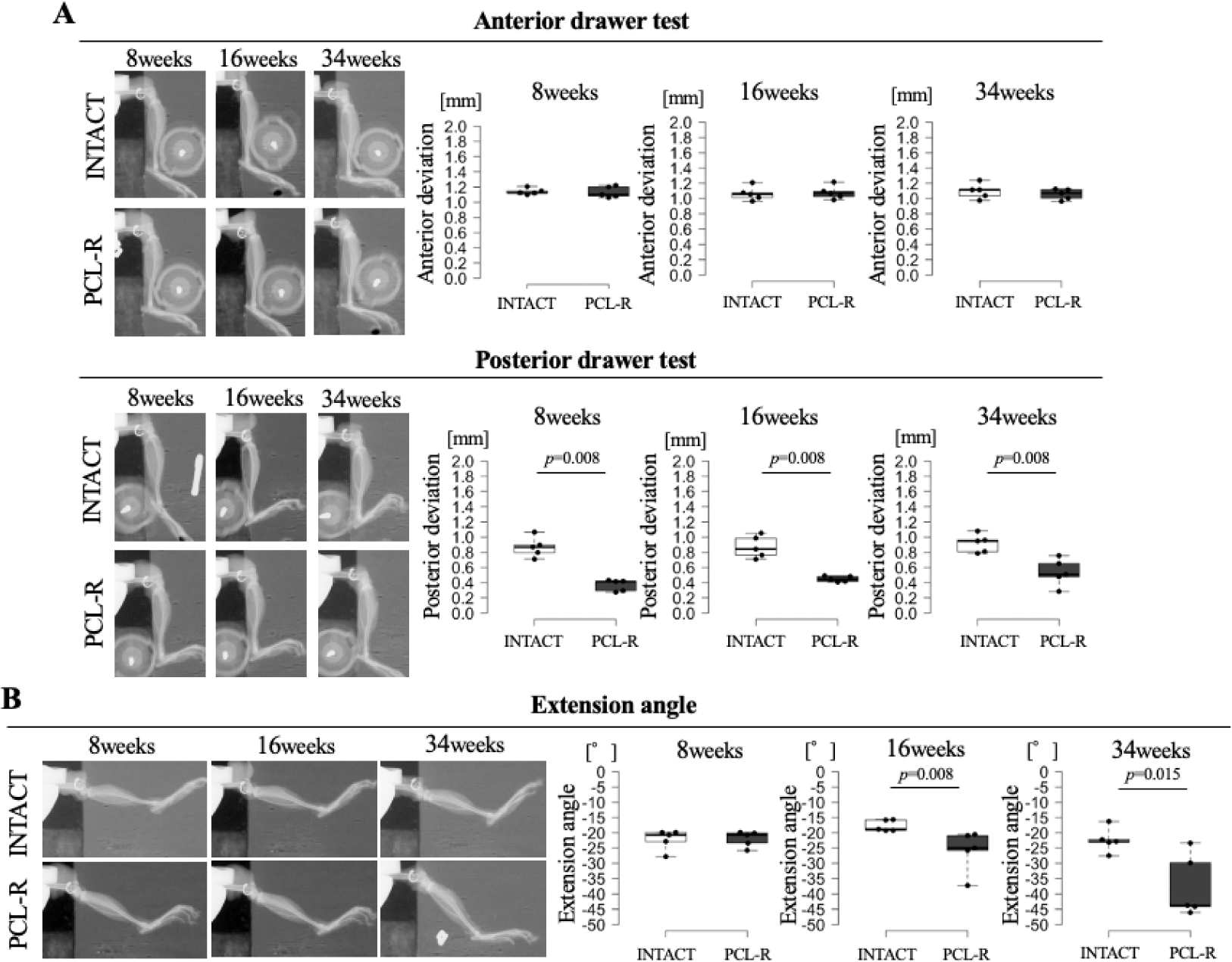
Evaluation of joint instability using soft x-ray analysis. (A) Results of the anterior drawer test. There were no significant difference between PCL-R and INTACT. Results of the posterior drawer test. At all timepoints, the amount of posterior tibial deviation in the PCL-R groups were significantly increased compared to that in the INTACT groups (p=0.008). (B) Results of the extension angle test. At 16 and 34 weeks, the extension angle in the PCL-R groups were significantly increased compared to that in the INTACT groups. All data are presented as the median with interquartile range.

### Observation of the ruptured PCL in the long term

The HE staining images are shown in Fig.5. At all timepoints, in the INTACT group, the continuity of the PCL was maintained, and the cells were arranged in a regular pattern, whereas in the PCL-R group, the continuity of the PCL was disrupted, and the cells were arranged in an irregular pattern.

**Figure 5.**
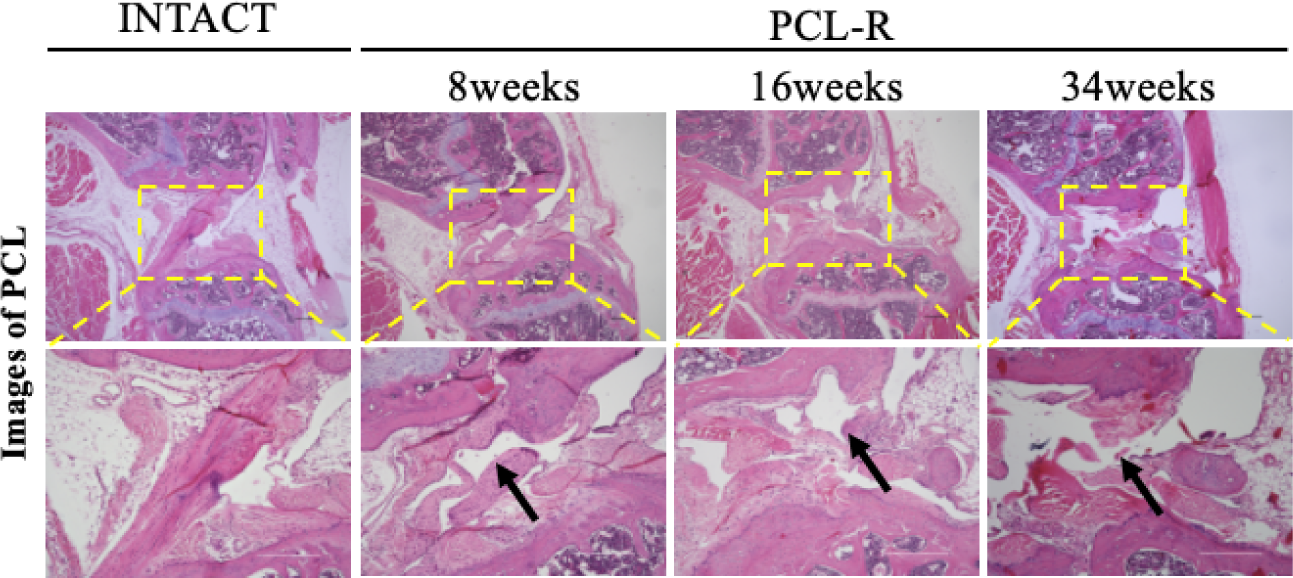
The images of the ruptured PCL. Unlike INTACT, where cells are arranged regularly, the arrangement of cells is disordered and continuity in PCL-R is lost. Furthermore, the loss of continuity in PCL-R lasted up to 34 weeks. Scale bar: 300 mm.

### PCL-R group showed slight cartilage degeneration compared to DMM group

The safranin-O/fast green staining images are shown in Fig.6(A). Regarding cartilage degeneration, it was observed in the central tibial region of the DMM group. In contrast, in the PCL-R group, cartilage degeneration was confirmed in the anterior tibial region. The OARSI score results are shown in Fig.6(B). At 8 weeks, there were no significant differences in OARSI score among the three groups (*p*=0.75). At 16 weeks, the OARSI score in the DMM group was significantly higher than that in the INTACT group (*p*=0.019). On the other hand, there was no significant difference in the OARSI scores of the PCL-R group compared with the other two groups (PCL-R vs INTACT; *p*=0.08, PCL-R vs DMM; *p*=0.12). At 34 weeks, the OARSI score in the DMM group was significantly higher than that in the INTACT group (*p*=0.024). Conversely, at 34 weeks, the PCL-R group showed no significant difference in OARSI score compared with the other two groups (PCL-R vs INTACT; *p*=0.35; PCL-R vs DMM; *p*=0.11).

**Figure 6.**
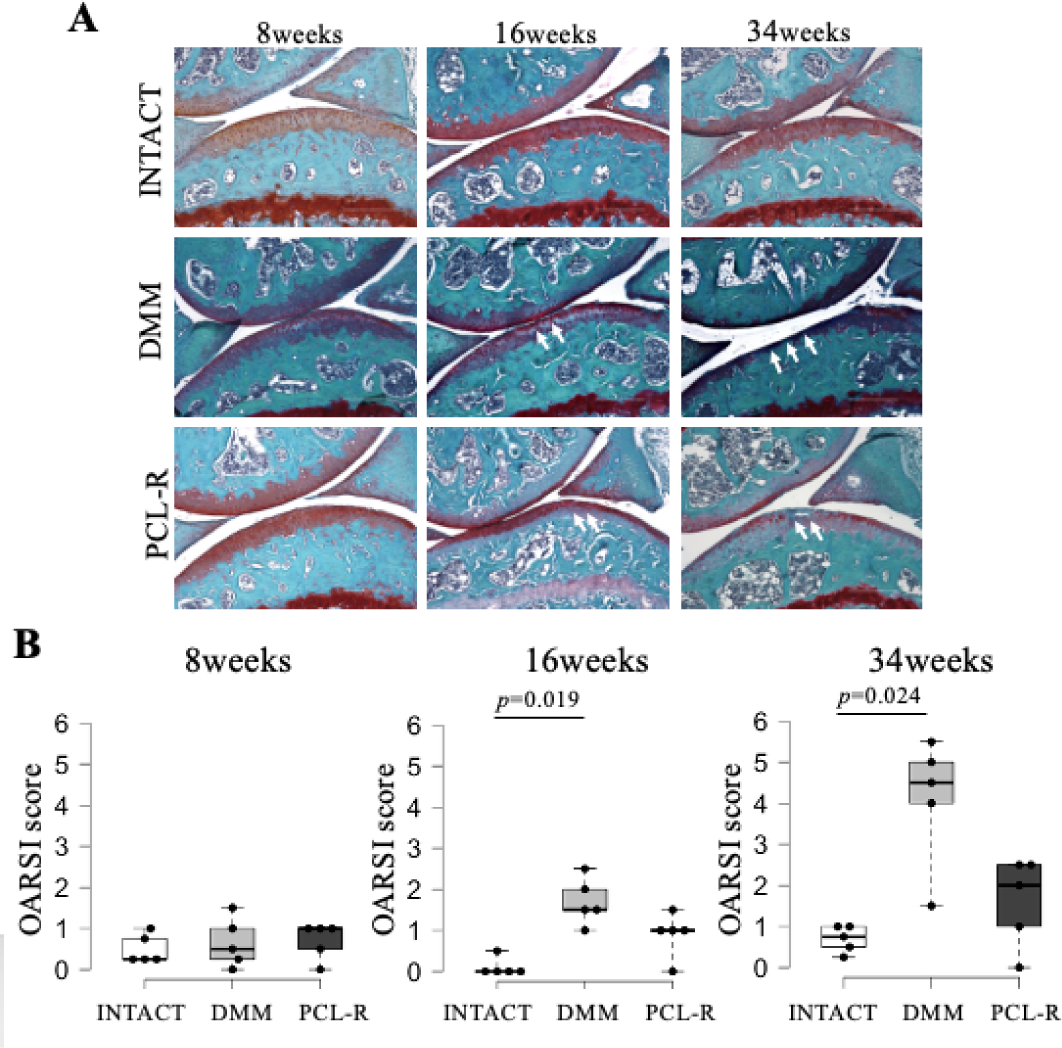
Evaluation of articular cartilage degeneration. (A) Representative safranin-O/fast green-stained image of each group. The histological images show articular cartilage of the medial tibia. In the PCL-T group, irregularity and fibrillation of articular cartilage were observed. Arrows indicate cartilage degeneration sites. Degeneration is localized in the central region in the DMM group and in the posterior region in the PCL-R group. (B) The results of OARSI score. DMM groups were significantly higher in OARSI scores than INTACT at 16 and 34weeks. The data are presented as the median with interquartile range. Scale bar: 300 mm.

The results of immunohistochemical staining of Col X are shown in Fig.7. At 8 weeks, Col X-positive cell ratio in the PCL-R group was significantly higher than that in the INTACT group (*p*=0.024). At 16 weeks, the Col X-positive cell ratio in the DMM group was significantly higher than that in the INTACT group(*p*=0.043). On the other hand, there was no significant difference in the Col X-positive cell ratio of the PCL-R group compared with the other two groups (PCL-R vs INTACT; *p*=0.12, PCL-R vs DMM; *p*=0.95). At 34 weeks, there were no significant differences among the three groups (*p*=0.13).

**Figure 7.**
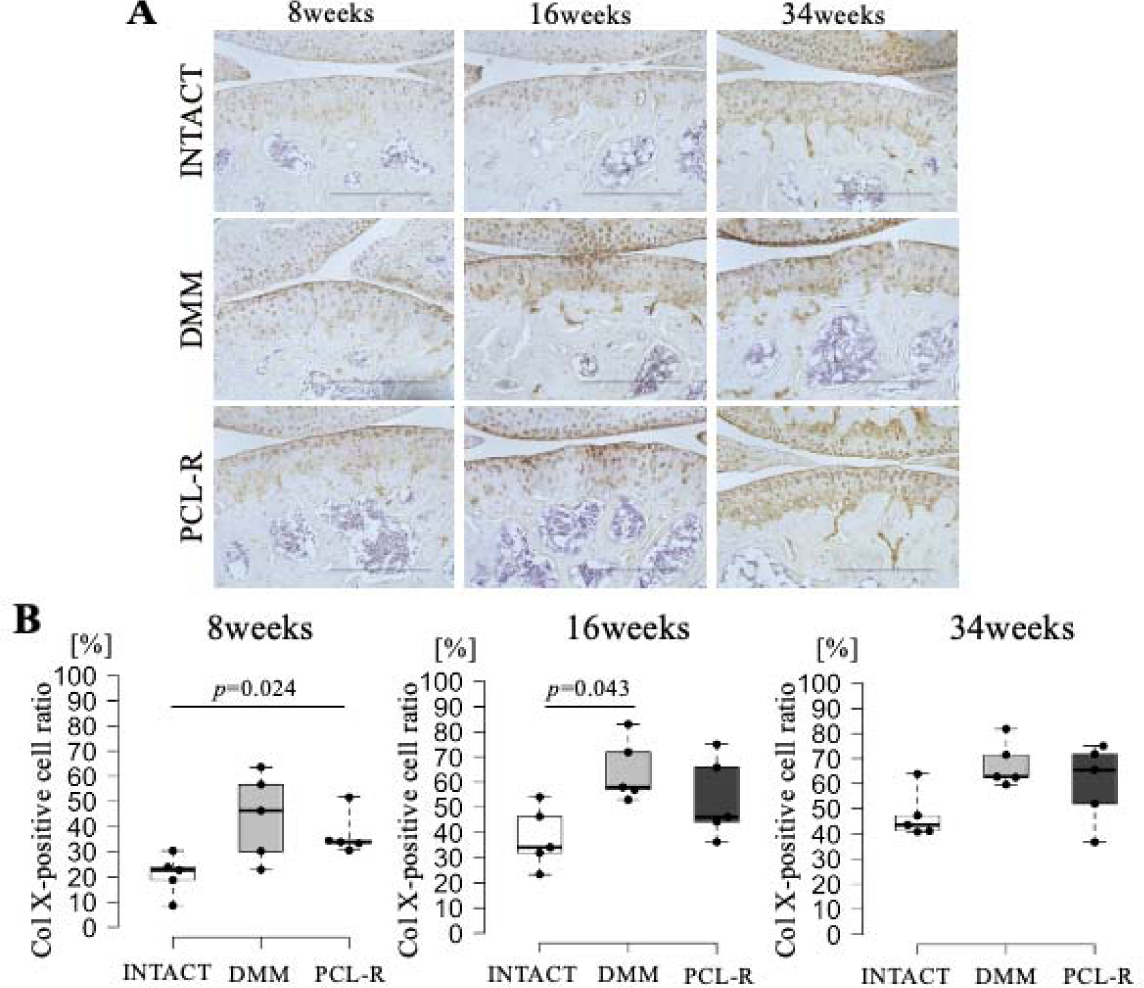
Comparison of Col X-positive cell rates between models. (A) Representative the IHC stained image of each group. (B) The results of Col X-positive cell ratio. The positive cell ratio in PCL-R was significantly higher than that in INTACT at 8 weeks (P=0.024). The positive cell ratio in DMM was significantly higher than that in INTACT (P=0.043) at 16 weeks. There were no significant differences among three groups at 34 weeks. The data are presented as the median with interquartile range. Scale bar: 300 mm.

### Subchondral bone structure in the PCL-R group showed no OA change

The reconstruction image of the medial side of the knee joint obtained by micro-CT analysis is shown in Fig.8 (A). The results of the subchondral bone construction were shown in Fig.8 (B). We calculated the *BV/TV*, *Tb.Th*, *Tb.N* and *Tb.Sp*. There were no significant differences among the three groups of the *BV/TV* at all timepoints (at 8 weeks; *p*=0.36, at 16 weeks; *p*=0.11, at 34 weeks; *p*=0.25). At 8 weeks, there were no significant differences among the three groups (*p*=0.99) of the *Tb.Th*. At 16 weeks, the *Tb.Th* in the DMM group was significantly higher than that in the INTACT group (*p*=0.043). However, there was no significant difference between the *Tb.Th* in the PCL-R group and that in the INTACT group (*p*=0.12). At 34 weeks, there were no significant differences among the three groups (*p*=0.18). At 8 and 16 weeks, there were no significant differences among the three groups of the *Tb.N* (8 weeks; *p*=0.68, 16 weeks; *p*=0.056). At 34 weeks, *Tb.N* in the PCL-R group was significantly higher than that in the DMM group (*p*=0.024). There were no significant differences among the three groups of he *Tb.Sp* at all timepoints (8 weeks; *p*=0.18, 16 weeks; *p*=0.21. 34 weeks; *p*=0.37).

**Figure 8.**
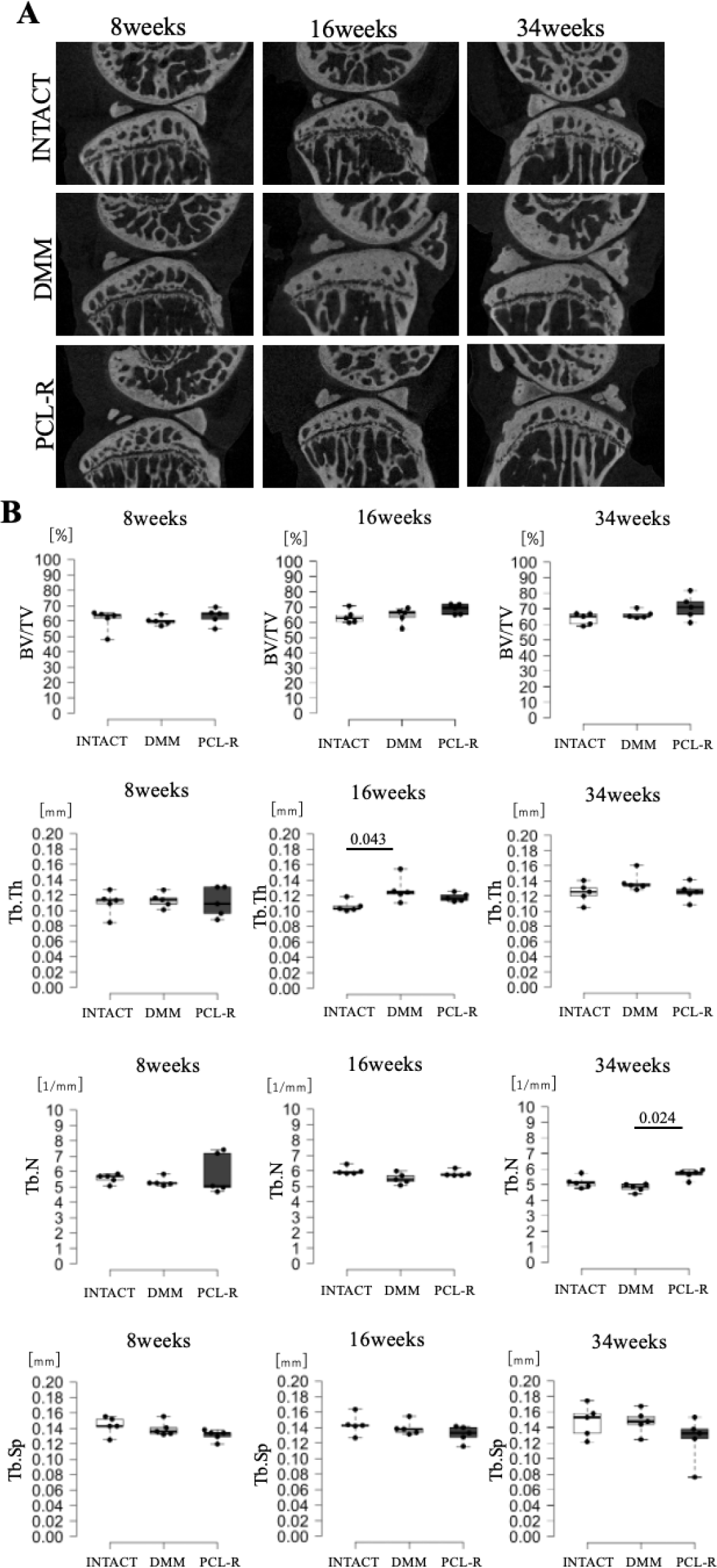
Results of the micro-CT analysis. (A) Representative image of the subchondral bone. (B) Results of the analysis of bone parameters using micro-CT. For BV/TV, no significant differences were observed among the three groups at any time point. For Tb.Th, no significant differences were found at 8 and 34 weeks, whereas Tb.Th in the DMM group was significantly higher than that in the INTACT group at 16 weeks (P = 0.043). For Tb.N, no significant differences were observed at 8 and 16 weeks, while Tb.N in the PCL-R group was significantly higher than that in the INTACT group at 34 weeks (P = 0.024). For Tb.Sp, no significant differences were detected among the three groups at any time point. Data are presented as medians with interquartile ranges.

## DISCUSSION

Knee OA is a multifactorial disease composed of cartilage and subchondral bone degeneration with aging. To elucidate this complex onset mechanism, alternatives to animal testing are emerging in knee OA research. Whereas animal models remain critical for understanding the onset mechanisms of joint degeneration, particularly in vivo tissue interactions. Here, we developed a novel PCL-R knee OA model with characteristics of non-invasiveness, non-intra-articular injury, and gradual cartilage degeneration over a longer time course than current knee OA models. This model was created by manually posteriorly dislocating the tibia, resulting in rupture of the PCL at the mid-parenchymal portion. The temporal changes in the severity of articular cartilage degeneration and subchondral bone in PCL-R models induced articular cartilage degeneration in a sequential and steady manner. Compared to conventional models, cartilage degeneration in PCL-R models progressed more slowly, demonstrating their effectiveness as a model for elucidating the onset mechanism of knee OA.

This novel PCL-R model is quite simple, reproducible, and practical, inducing posterior deviation solely by manually dislocating the tibia. The PCL rupture forces ranged from 8.56 to 12.7 N. The PCL rupture force was comparable to the ACL rupture force in the ACL-R model, as previously established ^20^. In addition, based on the results immediately after the PCL-R model was created, posterior displacement occurred in the PCL-R group, with no intra-articular microfractures or cartilage defects observed. Additionally, the PCL-R model is non-invasive and does not involve articular capsule resection. Surgically induced models, widely used in this field, certainly cause unnecessary acute inflammation^22^ and excessive abnormal joint motion following resection of surrounding tissues, potentially accelerating the progression of knee OA as a model-specific phenotype. Conversely, the novel PCL-R model, which has no unintended events on cartilage and surrounding tissues due to the developing intervention, may be a useful tool for investigating the mechanisms underlying the onset and progression of knee OA.

Articular cartilage degeneration is the primary pathological change in knee OA. In this study, histological images showed complete rupture in the central portion of the PCL. Then, the PCL-R group showed a significant increase in posterior joint deviation compared with the INTACT group at 8, 16, and 34 weeks. These results indicate that PCL rupture leads to sustained posterior joint deviation in the PCL-R group. Consequently, cartilage degeneration in the PCL-R group was confirmed in the anterior tibial region, involving posterior tibial deviation due to PCL rupture. However, there was no significant difference in OARSI score between the PCL-R and INTACT groups at 34 weeks post-surgery. On the other hand, cartilage degeneration was clearly observed in the central tibial region of the DMM group, and the OARSI score showed a significant difference compared with the INTACT group at 16 and 34 weeks after surgery. It has been reported that articular cartilage degeneration occurred depending on joint instability ^23^. In addition, we previously reported that suppressing joint instability inhibits cartilage degeneration^24^. Therefore, compared with the DMM group, the PCL-R group showed a lower impact of joint instability, but instability certainly occurred, suggesting that cartilage degeneration may have progressed more slowly. Interestingly, differences in the sites of cartilage degeneration were observed between the DMM and PCL-R groups. In particular, the PCL-R group showed cartilage degeneration in the anterior tibial region. This is thought to be caused by concentrating mechanical stress on the anterior tibia due to the posterior tibial deviation. This point is one of the limitations of the PCL-R model as a model-specific feature for discussing the onset mechanism of knee OA based on the data from this model.

The hypertrophy of articular surface chondrocytes labeled by Col-X represents an initiating stage of cartilage degeneration^25^. In particular, it has been demonstrated that Col-X-positive chondrocytes express MMP13 even before significant articular cartilage degeneration is confirmed by Safranin-O/fast green staining^25^. In this study, the OARSI scores in the PCL-R group were not significantly different from those in the INTACT group, whereas the Col-X positive cell ratio of PCL-R mice was significantly higher than that in the INTACT group at 8 weeks after surgery. On the other hand, the Col-X positive cell ratio in the DMM group was significantly higher than that in the INTACT group at 16 weeks after surgery. These results suggest that chondrocyte hypertrophy in the PCL-R model occurs earlier than in the DMM model and then gradually progresses to knee OA. It has been reported that mild to moderate degeneration occurs in DMM models within 4 to 8 weeks, and this model has been used as the most common OA model^14,24,26^; however, in current existing models, cartilage degeneration progresses within a few weeks after the intervention. Therefore, these models may overlook important factors in the pre- and onset of cartilage degeneration. The present study showed that the PCL-R models progress cartilage degeneration more slowly than the DMM model. Therefore, it can be inferred that the PCL-R model can thoroughly analyze the complex pathophysiology of knee OA without overlooking key factors in its onset and progression.

Subchondral bone changes are also one of the important characteristics of the early phase of knee OA. In knee osteoarthritis, bone resorption is accelerated in the early stages, while bone formation occurs in later stages^27^. Previous studies have reported that the features of current existing OA models, such as the DMM model and ACL-T model, exhibit subchondral bone changes in the early postoperative period^28–30^. Our results of micro-CT reconstruction images indicated that, compared with the INTACT group, the DMM group observed progressive subchondral bone formation from 16 weeks post-surgery onwards (Fig. 8A). In contrast, the PCLR group maintained the bone structure of the subchondral bone even at 34 weeks (Fig. 8A). In addition, there were no significant differences between the PCL-R group and the INTACT group for any bone parameters (Fig. 8B). These results suggest that changes in the subchondral bone were more gradual in the PCLR group than in the DMM group. However, in the analysis of bone parameters, the BV/TV in the DMM group was comparable to that in the INTACT group at all time points (Fig. 8B). Previous studies have reported that the features of current existing OA models, such as the DMM model and ACL-T model, exhibit subchondral bone changes in the early postoperative period^28–30^. In this study, no significant differences were observed between models in the analysis of bone parameters across the entire medial tibial subchondral bone. Therefore, in the future, it will be necessary to evaluate the dynamics of osteoclasts and osteoblasts in the subchondral bone and to assess the differences in subchondral bone remodeling among different models. In summary, we established the PCL-R model as a novel animal model for knee OA. The PCL-R model is used with non-invasive methods to rupture the PCL. In the PCL-R model, mild cartilage degeneration was observed at 16 to 34 weeks after surgery without involving any subchondral bone changes. The results of this study suggest that the PCL-R model exhibits a phenotype in which cartilage degeneration occurs more slowly than in conventional models. Furthermore, the gradual progression of cartilage degeneration more accurately reflects the pathophysiology of human knee OA. Therefore, the PCL-R model was suggested to be a potentially more useful model than conventional models for elucidating the onset mechanism of knee OA. A limitation of this study is that the contralateral limb of the PCL-R model was used as the INTACT group, and thus it was not a perfectly healthy knee joint. Therefore, it is necessary to analyze another sample as a pure INTACT group in the future. Furthermore, in this study, we performed only histological analysis of cartilage and structural analysis of subchondral bone. Knee OA should be considered as a whole-joint disease. Therefore, future studies should include detailed immunohistochemical and biochemical analyses of cartilage and meniscus, as well as of surrounding tissues such as synovium and synovial fluid.

In conclusion, this study established a novel PCL-R model in which OA develops more slowly than in conventional models. This study suggested that the PCL-R model is a useful model for elucidating the onset mechanism of knee OA.

## CONTRICUTIONS

All authors approved the final submitted manuscript.

Study design: SE, TK.

Mechanical analysis: SE, KT, RS, HM.

Data collection, Histological analysis: SE, KA, KT, YU, MS.

Digital microscope analysis: MS

Immunohistochemical analysis: KT

Statistical analysis: SE

Manuscript composition: SE, KA, TK.

## ACKNOWLEDGEMENTS

The author(s) received no financial support for the research, authorship, and/or publication of this article.

## REFERENCES

1. Hunter DJ, Bierma-Zeinstra S. Osteoarthritis. The Lancet. 2019;393(10182):1745–1759. doi:10.1016/S0140-6736(19)30417-9

2. Chen D, Shen J, Zhao W, et al. Osteoarthritis: toward a comprehensive understanding of pathological mechanism. Bone Res. 2017;5(1):16044. doi:10.1038/boneres.2016.44

3. Punzi L, Galozzi P, Luisetto R, et al. Post-traumatic arthritis: overview on pathogenic mechanisms and role of inflammation. RMD Open. 2016;2(2):e000279. doi:10.1136/rmdopen-2016-000279

4. Little CB, Hunter DJ. Post-traumatic osteoarthritis: From mouse models to clinical trials. Nature Reviews Rheumatology. 2013;9(8):485–497. doi:10.1038/nrrheum.2013.72

5. Liu Q, Niu J, Li H, et al. Knee Symptomatic Osteoarthritis, Walking Disability, NSAIDs Use and All-cause Mortality: Population-based Wuchuan Osteoarthritis Study. Scientific Reports. 2017;7(1):1–7. doi:10.1038/s41598-017-03110-3

6. Wang SX, Ganguli AX, Bodhani A, Medema JK, Reichmann WM, Macaulay D. Healthcare resource utilization and costs by age and joint location among osteoarthritis patients in a privately insured population. Journal of Medical Economics. 2017;20(12):1299–1306. doi:10.1080/13696998.2017.1377717

7. Kassebaum NJ, Arora M, Barber RM, et al. Global, regional, and national disability-adjusted life-years (DALYs) for 315 diseases and injuries and healthy life expectancy (HALE), 1990–2015: a systematic analysis for the Global Burden of Disease Study 2015. The Lancet. 2016;388(10053):1603–1658. doi:10.1016/S0140-6736(16)31460-X

8. De Luna A, Otahal A, Nehrer S. Mesenchymal Stromal Cell-Derived Extracellular Vesicles – Silver Linings for Cartilage Regeneration? Frontiers in Cell and Developmental Biology. 2020;8(December). doi:10.3389/fcell.2020.593386

9. Nedunchezhiyan U, Varughese I, Sun ARJ, Wu X, Crawford R, Prasadam I. Obesity, Inflammation, and Immune System in Osteoarthritis. Frontiers in Immunology. 2022;13(July):1–19. doi:10.3389/fimmu.2022.907750

10. Bapat S, Hubbard D, Munjal A, Hunter M, Fulzele S. Pros and cons of mouse models for studying osteoarthritis. Clinical & Translational Med. 2018;7(1):e36. doi:10.1186/s40169-018-0215-4

11. Little CB, Hunter DJ. Post-traumatic osteoarthritis: from mouse models to clinical trials. Nat Rev Rheumatol. 2013;9(8):485–497. doi:10.1038/nrrheum.2013.72

12. Lampropoulou-Adamidou K, Lelovas P, Karadimas EV, et al. Useful animal models for the research of osteoarthritis. Eur J Orthop Surg Traumatol. 2014;24(3):263–271. doi:10.1007/s00590-013-1205-2

13. Glasson SS, Blanchet TJ, Morris EA. The surgical destabilization of the medial meniscus (DMM) model of osteoarthritis in the 129/SvEv mouse. Osteoarthritis and Cartilage. 2007;15(9):1061–1069. doi:10.1016/j.joca.2007.03.006

14. SS G, Blanchet T, Morris E. The surgical destabilization of the medial meniscus (DMM) model of osteoarthritis in the 129 / SvEv mouse. Published online 2007:1061–1069. doi:10.1016/j.joca.2007.03.006

15. Ajalik RE, Linares I, Alenchery RG, et al. Human Tendon□on□a□Chip for Modeling the Myofibroblast Microenvironment in Peritendinous Fibrosis. Adv Healthcare Materials. 2025;14(4):2403116. doi:10.1002/adhm.202403116

16. Labusca L. Next-generation osteoarthritis models: integrating biological, computational, and engineering approaches. Stem Cell Res Ther. 2025;16(1):686. doi:10.1186/s13287-025-04790-9

17. Amis AA, Gupte CM, Bull AMJ, Edwards A. Anatomy of the posterior cruciate ligament and the meniscofemoral ligaments. Knee Surgery, Sports Traumatology, Arthroscopy. 2006;14(3):257–263. doi:10.1007/s00167-005-0686-x

18. Shelbourne KD, Clark M, Gray T. Minimum 10-year follow-up of patients after an acute, isolated posterior cruciate ligament injury treated nonoperatively. American Journal of Sports Medicine. 2013;41(7):1526–1533. doi:10.1177/0363546513486771

19. Christiansen BA, Anderson MJ, Lee CA, Williams JC, Yik JHN, Haudenschild DR. Musculoskeletal changes following non-invasive knee injury using a novel mouse model of post-traumatic osteoarthritis. Osteoarthritis and Cartilage. 2012;20(7):773–782. doi:10.1016/j.joca.2012.04.014

20. Takahata K, Arakawa K, Enomoto S, et al. Joint instability causes catabolic enzyme production in chondrocytes prior to synovial cells in novel non-invasive ACL ruptured mouse model. Osteoarthritis and Cartilage. 2023;31(5):576–587. doi:10.1016/j.joca.2022.12.004

21. Glasson SS, Chambers MG, Van Den Berg WB, Little CB. The OARSI histopathology initiative - recommendations for histological assessments of osteoarthritis in the mouse. Osteoarthritis and Cartilage. 2010;18(SUPPL. 3):S17–S23. doi:10.1016/j.joca.2010.05.025

22. Christiansen BA, Guilak F, Lockwood KA, et al. Non-invasive mouse models of post-traumatic osteoarthritis. Osteoarthritis and Cartilage. 2015;23(10):1627–1638. doi:10.1016/j.joca.2015.05.009

23. Kamekura S, Hoshi K, Shimoaka T, et al. Osteoarthritis development in novel experimental mouse models induced by knee joint instability. Osteoarthritis and Cartilage. 2005;13(7):632–641. doi:10.1016/j.joca.2005.03.004

24. Arakawa K, Takahata K, Enomoto S, et al. The difference in joint instability affects the onset of cartilage degeneration or subchondral bone changes. Osteoarthritis and Cartilage. 2022;30(3):451–460. doi:10.1016/j.joca.2021.12.002

25. Kamekura S, Hoshi K, Shimoaka T, et al. Osteoarthritis development in novel experimental mouse models induced by knee joint instability. Osteoarthritis and Cartilage. 2005;13(7):632–641. doi:10.1016/j.joca.2005.03.004

26. Hwang HS, Park IY, Hong JI, Kim JR, Kim HA. Comparison of joint degeneration and pain in male and female mice in DMM model of osteoarthritis. Osteoarthritis and Cartilage. 2021;29(5):728–738. doi:10.1016/j.joca.2021.02.007

27. Yuan XL, Meng HY, Wang YC, et al. Bone–cartilage interface crosstalk in osteoarthritis: potential pathways and future therapeutic strategies. Osteoarthritis and Cartilage. 2014;22(8):1077–1089. doi:10.1016/j.joca.2014.05.023

28. Sulaiman SZS, Tan WM, Radzi R, et al. Comparison of bone and articular cartilage changes in osteoarthritis: a micro-computed tomography and histological study of surgically and chemically induced osteoarthritic rabbit models. J Orthop Surg Res. 2021;16(1):663. doi:10.1186/s13018-021-02781-z

29. Iijima H, Aoyama T, Ito A, et al. Exercise intervention increases expression of bone morphogenetic proteins and prevents the progression of cartilage-subchondral bone lesions in a post-traumatic rat knee model. Osteoarthritis and Cartilage. 2016;24(6):1092–1102. doi:10.1016/j.joca.2016.01.006

30. Fang H, Huang L, Welch I, et al. Early Changes of Articular Cartilage and Subchondral Bone in The DMM Mouse Model of Osteoarthritis. Sci Rep. 2018;8(1):2855. doi:10.1038/s41598-018-21184-5

